# Therapeutic DBS for OCD Suppresses the Default Mode Network

**DOI:** 10.1101/2024.07.21.601827

**Authors:** Natalya Slepneva, Genevieve Basich-Pease, Lee Reid, Adam C. Frank, Tenzin Norbu, Andrew D Krystal, Leo P Sugrue, Julian C Motzkin, Paul S. Larson, Philip A. Starr, Melanie A. Morrison, A Moses Lee

## Abstract

**Background:** Deep brain stimulation (DBS) of the anterior limb of the internal capsule (ALIC) is an emerging treatment for severe, refractory obsessive-compulsive disorder (OCD). The therapeutic effects of DBS are hypothesized to be mediated by direct modulation of a distributed cortico-striato-thalmo-cortical network underlying OCD symptoms. However, the exact underlying mechanism by which DBS exerts its therapeutic effects still remains unclear.

**Method:** In five participants receiving DBS for severe, refractory OCD (3 responders, 2 non-responders), we conducted a DBS On/Off cycling paradigm during the acquisition of functional MRI to determine the network effects of stimulation across a variety of bipolar configurations. We also performed tractography using diffusion-weighted imaging (DWI) to relate the functional impact of DBS to the underlying structural connectivity between active stimulation contacts and functional brain networks.

**Results:** We found that therapeutic DBS had a distributed effect, suppressing BOLD activity within regions such as the orbitofrontal cortex, dorsomedial prefrontal cortex, and subthalamic nuclei compared to non-therapeutic configurations. Many of the regions suppressed by therapeutic DBS were components of the default mode network (DMN). Moreover, the estimated stimulation field from the therapeutic configurations exhibited significant structural connectivity to core nodes of the DMN.

**Conclusions:** Therapeutic DBS for OCD suppresses BOLD activity within a distributed set of regions within the DMN relative to non-therapeutic configurations. We propose that these effects may be mediated by interruption of communication through structural white matter connections surrounding the DBS active contacts.

## Introduction

Obsessive-Compulsive Disorder (OCD) is a common psychiatric disorder characterized by intrusive anxiety-provoking thoughts and repetitive behaviors. The symptoms of OCD are thought to result from aberrant activity within a cortico-striato-thalamo-cortical (CSTC) network involving the orbitofrontal cortex (OFC), medial prefrontal cortex (PFC), and interconnected basal ganglia and associated thalamo-cortical circuits [1–3]. Evidence-based treatments for OCD include cognitive behavioral therapy and medications, such as serotonin reuptake inhibitors [4]. However, it has been estimated that approximately 10% of patients continue to have severe, debilitating symptoms that are not addressed by conventional therapies [5].

Deep brain stimulation (DBS) is an invasive form of neuromodulation that has been used to treat severe cases of OCD [6–8]. DBS involves direct electrical stimulation delivered through electrodes implanted in deep structures in the brain to modulate neural circuits. The most common DBS target for OCD is the anterior limb of the internal capsule (ALIC), which receives topographically organized connections from various components of the CSTC network [9–12].

However, the underlying mechanism by which DBS mediates its therapeutic effects remains unclear. This lack of mechanistic understanding is a barrier to addressing two important clinical limitations of the treatment: 1) Only 60% of patients respond to DBS at the ALIC target [7, 8], and 2) DBS programming to find the optimal configuration of stimulation contacts and parameters currently involves a complex, trial-and-error process guided by inconsistent subjective reports that can take months to years. For this reason, there is a need to identify biomarkers of target engagement that are tied to therapeutic efficacy. Such a biomarker could be used to predict treatment response and guide more efficient DBS programming.

Advances in DBS technology now allow for the safe acquisition of magnetic resonance imaging (MRI) data while DBS is cycled On and Off. With these advances, we can now map the network impact of DBS on whole brain activity using functional MRI (fMRI) blood-oxygen-level-dependent (BOLD) imaging. The aim of this study was to determine the network effects of ALIC DBS by conducting fMRI while DBS was cycled On and Off in patients receiving DBS for severe, refractory OCD. We hypothesized that a common network related to OCD symptoms would be engaged specifically by therapeutic DBS configurations, and further, that our estimated therapeutic DBS stimulation fields would exhibit strong structural connectivity with this network.

## Methods and Materials

### Subject Recruitment

Five patients treated with DBS for their severe, refractory OCD were identified from the UCSF OCD Clinic. In all patients the DBS devices had been implanted under the FDA Humanitarian Device Exemption with bilateral leads targeting to the anterior limb of the internal capsule (ALIC) and neighboring bed nucleus of the stria terminalis (BNST). At the time of the study, all patients had a Medtronic Percept PC DBS stimulator (Medtronic, Minneapolis, MN), which is MRI conditional at 3 Tesla (3T). Institutional review board approval and MRI safety committee approval was obtained prior to initiating recruitment. All subjects provided written informed consent for the study.

Three of the five subjects demonstrated marked clinical improvement of their OCD symptoms in response to DBS and were classified as treatment responders, defined as a greater than 35% reduction in their last Yale-Brown Obsessive Compulsive Severity (YBOCS) compared to their pre-surgical baseline. The remaining two subjects exhibited minimal clinical improvement and were classified as treatment non-responders. In these subjects more than a year was spent trying to identify therapeutic settings without success, and their DBS devices are currently off (Table 1).

### DBS Cycling Paradigm

We acquired 6 minute runs of fMRI data at 3T using a block design where the DBS device was cycled On and Off for one-minute intervals including 8 seconds to ramp stimulation up and down at the beginning of the DBS On and Off block, respectively. The Medtronic DBS Percept PC device is MR Conditional only in the bipolar configuration. During each fMRI run, stimulation was delivered in a bipolar configuration at a pair of adjacent electrode contacts on either the left or right brain lead. For each run the active contact pair was chosen randomly from 12 possible configurations. Configurations were tested outside the scanner for tolerability before being trialed within the scanner. If configurations were not tolerable, they were not tested in the scanner. Due to time constraints, only a subset of these configurations was tested in each subject. At the beginning of each scanning session the DBS device was set in cycling mode, and the fMRI runs were timed to begin at the start of the Off cycle. Stimulation amplitude was set at the maximum cycling amplitude tolerated, which was either 5 or 6mA for all patients.

The Medtronic Percept device is only labeled for MR imaging with bipolar stimulation. However, in treatment responders, the active treatment electrode configuration could be either a bipolar or monopolar configuration. For our fMRI experiments if the responder’s active configuration was a bipolar setting this was considered the therapeutic configuration for that electrode. If the responder’s active configuration was a monopolar setting, then the therapeutic configuration for that electrode was defined as the bipolar configuration for that electrode that shared the same anode as the active monopolar setting.

### Imaging Acquisition

MRI scans were acquired on a 3T whole body scanner in low specific absorption rate mode (Discovery MR750, GE Healthcare, Chicago, IL) with a 32-channel receive head coil (Nova Medical). For all subjects, multiple runs of gradient-echo fMRI data were acquired with the following parameters: TR/TE=2s/30ms, voxel size=3.75×3.75×4 mm, flip angle=86 degrees, FOV=240mm, in-plane acceleration factor=2, run length=6 minutes. Manufacturer guidelines, allow a maximum of 30 min of MRI scanning with the Medtronic Percept during a single visit. Thus during each visit multiple 6 min runs testing different configurations were acquired up to a total of 30 min of scan time. Between runs, the participant was removed from the scanner to an MRI safe zone while lying on the detached scanner bed with their head in the same position so the study clinician could change the DBS settings before being returned to the MRI. Some participants returned for multiple visits over separate days to acquire additional scans with different configurations.

For three subjects (one responder, two non-responders), we collected pre-surgical diffusion weighted images (DWI) (3T, 55 direction, b=2000 s/mm^2^). For two of the subjects (both responders), no pre-surgical DWI was available, so we collected post-implant T1-weighted (T1w) structural MRI and DWI (3T, 29-direction, b=1000 s/mm^2^) for these participants. We have previously demonstrated that tractography using post-implant DWI is feasible and reproduces results from pre-implant DWI [13].

### Image Preprocessing and Denoising

T1 and fMRI data preprocessing was performed with fMRIPrep 21.0.1 [14], a standard preprocessing pipeline based on Nipype 1.6.1[15], which uses Nilearn 0.8.1 [16] for many internal operations. Anatomical preprocessing generated a subject-specific T1w reference template for registration of fMRI images to T1w (subject) and MNI spaces. In brief, the anatomical preprocessing steps included the following: T1w images from all scanning sessions for an individual subject were bias field corrected with N4BiasFieldCorrection [16] in ANTs 2.3.3 [17] and averaged to generate a subject-specific anatomical reference image using mri_robust_template (FreeSurfer 6.0.1) [18]. The T1w reference was skull-stripped using FAST (FSL 6.0.5.1) [19] then normalized to the MNI152NLin6Asym standard space via nonlinear registration with ANTs.

For each fMRI run, skull-stripped and non-skull-stripped BOLD reference volumes were co-registered to subject (T1w) and MNI space. Head-motion parameters with respect to the fMRI reference (transformation matrices and six corresponding rotation and translation parameters) were estimated before any spatiotemporal filtering using mcflirt (FSL 6.0.5.1) [20]. BOLD runs were slice-time corrected to 0.98s (mean of slice acquisition range 0s-1.96s) using 3dTshift from AFNI [21, 22]. The slice-time-corrected BOLD time-series were resampled onto their original, native space by applying the transforms to correct for head-motion. The BOLD reference was then co-registered with six degrees of freedom to the T1w reference using bbregister (FreeSurfer) which implements boundary-based registration [23]. Framewise displacement (FD) was calculated based on the preprocessed BOLD using two formulations, absolute sum of relative motions [24] and relative root mean square displacement between affines [25], using Nipype [15]. Frames that exceeded a threshold of 0.5 mm FD or 1.5 standardized spatial standard deviation of successive frames (DVARS) were annotated as motion outliers. The BOLD time-series were then resampled into standard space using Lanczos interpolation in antsApplyTransforms, generating a preprocessed BOLD run in MNI152NLin6Asym space. All the pertinent transformations (i.e. head-motion transform matrices and co-registrations to anatomical and output spaces) were composed into a single resampling step. Independent component analysis (ICA) using FSL MELODIC [26] was performed on the preprocessed BOLD in MNI space time-series after removal of non-steady state volumes and spatial smoothing with an isotropic, Gaussian kernel of 6mm FWHM. Noise components were identified using ICA-AROMA [27].

Preprocessed fMRI data were reviewed for quality, and ICA-derived noise components were manually identified by three expert raters (N.S. & M.A.M & G.B.P). ICs identified as noise by both raters were removed using MELODIC to generate an ICA-denoised BOLD timeseries. We removed initial non-steady state volumes from the denoised BOLD timeseries, then scaled the timeseries to a mean of 100.

### Image Post-processing and Analysis

To generate activation maps associated with DBS cycling for each subject, we used 3dDeconvolve in AFNI to perform ordinary least squares multiple regression on the preprocessed and denoised fMRI data in MNI space. Each model included a boxcar regressor convolved with a standard hemodynamic response function corresponding to On and Off periods, as well as nuisance regressors for 3 polynomial drift terms. High motion timepoints (i.e., FD >0.5 or DVARS > 1.5) were censored from analysis. Resulting parameter estimates corresponding to On and Off regressors were used to generate [On] – [Off] contrast maps for each individual run.

Group comparisons were performed using a 3dLME, a linear mixed-effects (LME) model in AFNI, to compare On-Off changes in BOLD signal between therapeutic and nontherapeutic stimulation configurations. In the LME model, the responder status and therapeutic configurations were modeled as fixed effects and subjects were modeled as random effects.

### DWI processing and Electrode Reconstruction

Preprocessing was performed using standard methods in QSIPrep 0.15.2, which is based on Nipype 1.7.0 [15]. DWI preprocessing in QSIPrep included the following steps: using MRtrix3, MP-PCA denoising was applied to DWI volumes with dwidenoise [28], Gibbs unringing was performed using mrdegibbs [32], and B1 field inhomogeneity was corrected with dwibiascorrect using the N4 algorithm [29]. Eddy current and head motion correction were performed using eddy (FSL 6.0.5.1) [34], and eddy’s outlier replacement was run with default parameters. Finally, DWI time-series were resampled to the ACPC coordinate system and 1mm isotropic voxels using the Jacobian modulation interpolation method in eddy. [28–31]

We used Lead-DBS 2.6 [32], a MATLAB toolbox for DBS electrode reconstruction and simulation of DBS, to model diffusion tracts stimulated by DBS. First, individual pre- and postoperative T1w, T2w (where available), and DWI scans were co-registered using SPM12 [17] and normalized to MNI152NLin2009b space using ANTs. Post-operative T1w anatomical scans were used to manually localize DBS electrode locations. DBS electrodes were then reconstructed for each subject. Using FastField [37], which utilizes a volume conductor model of the DBS electrode and surrounding tissue, the volume of activated tissue (VAT) by stimulation was modeled for each bipolar stimulation configuration that was used during fMRI runs. We generated individualized whole-brain tractography from denoised diffusion imaging data using the DSI studio implementation of the generalized q-sampling [33] imaging method (GQI) in Lead Connectome. 200,000 fibers were estimated for each subject, using default parameters for the GQI tracking [34]. For each patient and bipolar stimulation configuration, whole-brain tractograms were filtered to isolate streamlines that passed through the VAT. These remaining streamlines were used to estimate connectivity to parcels from Schaefer cortical [35] and Tian subcortical atlases [36].

### Statistics for Network Analysis

The Schaefer 100-parcel, seven network fMRI atlas was used to estimate the effect of the DBS On-Off response across these canonical networks for each individual fMRI run. We utilized a one-sided bootstrap spatial permutation test to determine if there was a significantly increased BOLD suppression within one of the seven networks for the therapeutic and non-therapeutic runs. To do this, we calculated the observed average amount of suppression across parcels within each of the seven networks for the therapeutic and non-therapeutic runs. We then generated a surrogate distribution (n=10,000) derived by permuting the parcels assigned to each of the seven networks to separately calculate the average amount of suppression within each network for the therapeutic runs and non-therapeutic runs. P-values were then calculated by finding the proportion of surrogates with suppressions greater than the actual differences observed, yielding one-tailed p-values for each network. Likewise, a similar two-sided bootstrap method was used to calculate whether there was a significant difference in BOLD activation/suppression within each of the seven networks between the therapeutic and non-therapeutic configurations. A Bonferroni correction was used to correct for multiple comparisons across the seven networks.

A similar approach using the Schaefer 100 parcel-7 network fMRI atlas was used to quantify the structural connectivity as measured by the fraction of total streamlines from the estimated VAT to cortical parcels within the canonical networks. A similar one-sided spatial bootstrap test was used to determine if there was a significantly increased total number of streamlines to one of the seven networks for the therapeutic configurations across responders and non-therapeutic configurations across all subjects. This method was also used to determine if there was a significant difference in fraction of total streamlines between therapeutic and non-therapeutic configurations in the seven networks.

## Results

Five patients who had been treated with DBS for their severe, refractory OCD were identified from the UCSF OCD Clinic. Their DBS leads were located within the ALIC and neighboring BNST (Fig 1A). For each of the patients, structural and diffusion-weighted MRI data were collected to identify the structural connectivity from the estimated volume of activated tissues (VAT) from different electrode stimulation configurations (Fig 1B). fMRI data were acquired at 3 Tesla while the DBS device was set in a one-minute ON/one-minute OFF cycling paradigm (Fig 1C). Subsequently, DBS On-vs -Off contrast maps were generated for each DBS configuration (Fig 1D). Group contrast maps between DBS On vs Off for the therapeutic (Supplemental Fig S1) and non-therapeutic (Supplemental Fig S2) configurations were derived. For therapeutic DBS configurations, we identified significant suppression in components of the CSTC circuit implicated in OCD such as the right orbitofrontal cortex, bilateral dorsomedial prefrontal cortex, left subthalamic nuclei, and right thalamus as well as other regions such as the precuneus and posterior cingulate cortex (Supplemental Fig S1, p<0.05 LME). For non-therapeutic DBS configurations, we found heterogenous changes, which were often not consistent across configurations or participants (Supplemental Fig S2, p<0.05 LME). We also generated a difference map between the therapeutic vs non-therapeutic configurations for DBS On-vs-Off, which identified significant BOLD suppression in the right orbitofrontal cortex, bilateral dorsomedial prefrontal cortex, and bilateral subthalamic nuclei, components of the CSTC network, as well as the left dorsolateral prefrontal cortex, precuneus, and posterior cingulate cortex (Fig 2, p<0.05, LME). In general, suppressions of BOLD activity for therapeutic configurations with DBS On-versus-Off, and the difference map between therapeutic and non-therapeutic configurations, corresponded to regions distant from the sites of the active electrode contacts located in the ALIC. However, it is possible that local BOLD signal changes within the ALIC itself may have been obscured by the presence of the electrode artifact.

**Figure 1:**
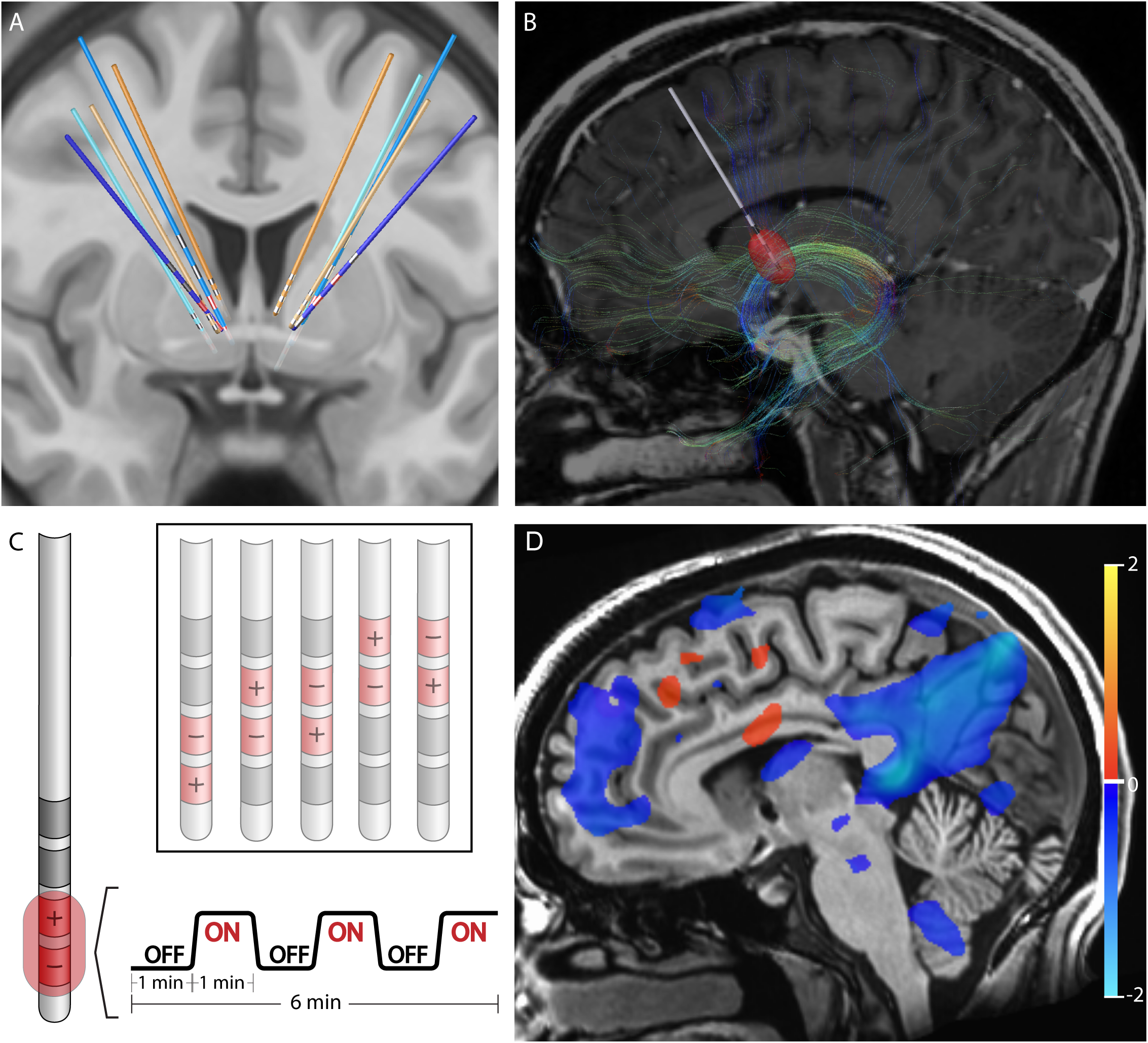
Structural and Functional Characterization of DBS Configurations. A) Reconstruction of DBS leads for five subjects. Leads for separate patients are in distinct colors. Blue leads indicate treatment responders; orange leads indicate non-responders. Therapeutic electrode contacts shown in red. B) Example of tractography derived from diffusion imaging seeding the estimated volume of tissue activation for a therapeutic bipolar contact configuration for a single representative subject. C) Design of DBS cycling On vs Off paradigm during fMRI acquisition for different stimulation configurations. D) fMRI BOLD changes with DBS On vs Off for same subject and configuration in B. Suppression of BOLD with DBS On vs Off is depicted in blue while activation with DBS On vs Off is depicted in red. Color bar indicates percentage change in BOLD signal. p<0.05; OLSQ.

**Figure 2.**
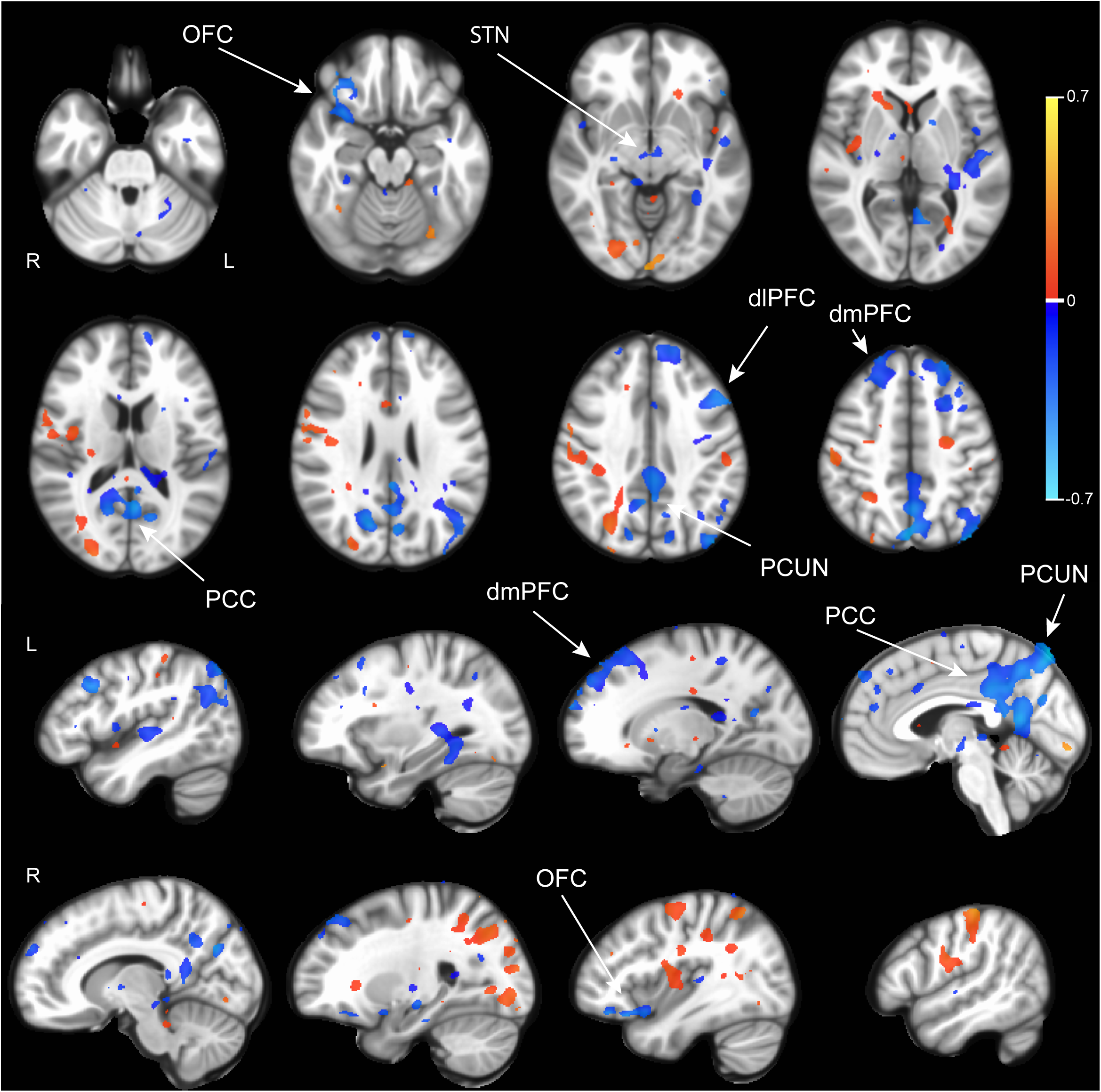
BOLD response differences between therapeutic and non-therapeutic DBS. Group comparison using linear mixed effects model of BOLD response between DBS On and Off in therapeutic (n=6 runs, 3 subjects) vs nontherapeutic (n=17 runs, 5 subjects) DBS configurations. Activations are in red and suppressions in blue. Color bar indicates percentage change in BOLD signal. dmPFC: dorsomedial prefrontal cortex. OFC: orbitofrontal cortex. PCUN: precuneus. PCC: posterior cingulate cortex. STN: subthalamic nucleus. p<0.05; LME.

Next, we asked whether the observed pattern of DBS On-versus-Off BOLD responses localized to particular functional networks by extracting network-specific parameter estimates for each contrast using existing network atlases (Supplemental Fig S3). Comparing DBS On-vs-Off for therapeutic configurations revealed a significant decrease in BOLD signal within the default mode network (DMN) (p= 1.0×10^-4^, one-sided permutation test with Bonferroni correction). We also observed a significant difference in BOLD signal change between therapeutic and nontherapeutic DBS On-vs-Off in the DMN (p=1.0×10^-4^, one-sided permutation test with Bonferroni correction) and control network (p=2.2×10^-3^, one-sided permutation test with Bonferroni correction) (Fig 3).

**Figure 3.**
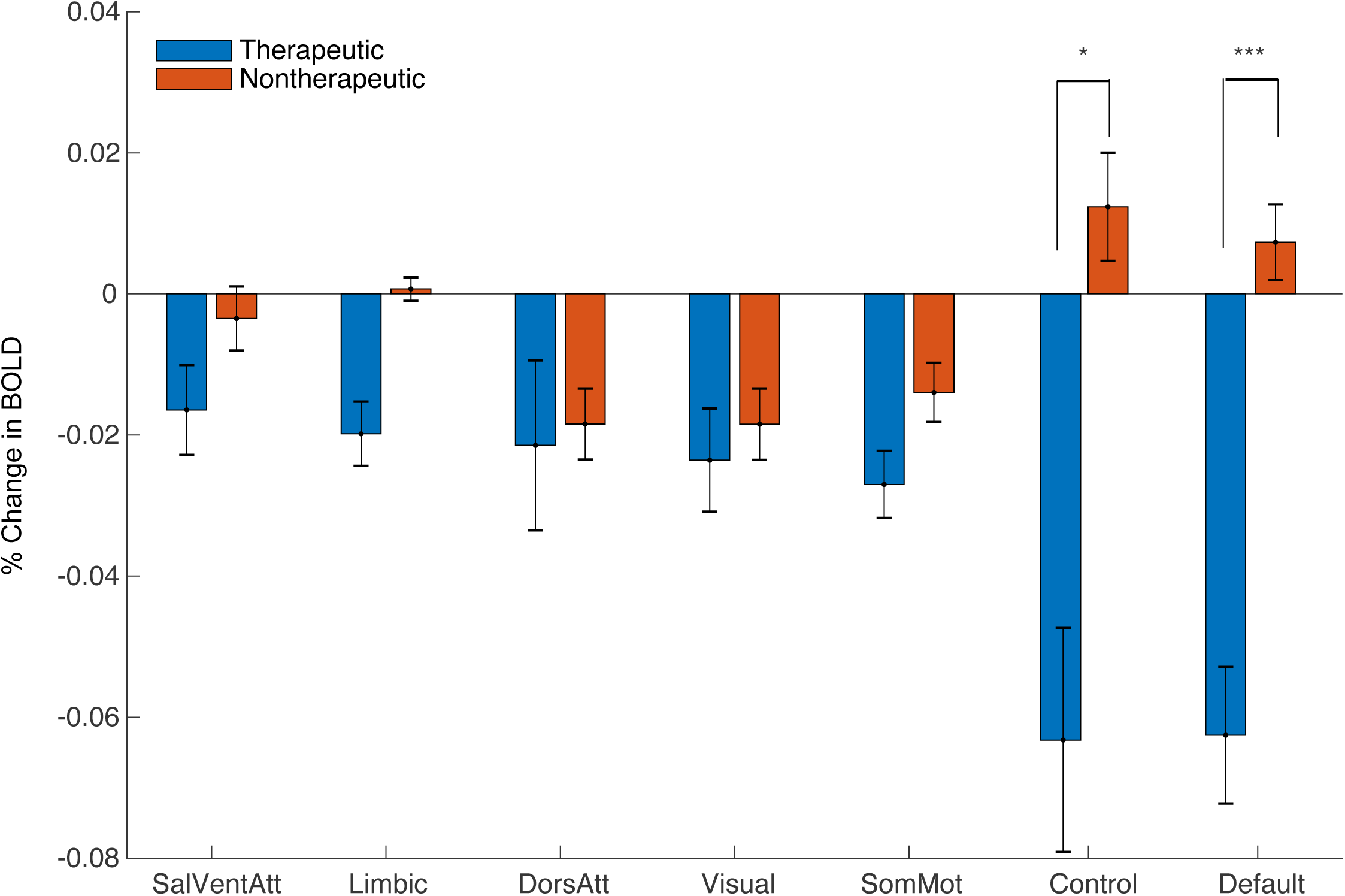
Network Impact of Therapeutic and Non-Therapeutic DBS. Comparison of average BOLD changes within canonical resting-state networks for therapeutic (blue) and non-therapeutic (red) configurations. ***p<5.0×10^-3^, *p<.05; permutation test (one-sided with Bonferroni correction).

Finally, we sought to determine whether structural connectivity from the estimated VAT for each DBS configuration identified similar functional networks by comparing the fraction of streamlines reaching each network parcel (Supplemental Fig S4). For the therapeutic configurations, we found significantly increased fraction of structural connections to the DMN relative to other networks (Fig 4A,B; p=0.011, one-sided permutation test). We also found that therapeutic configurations had significantly more structural connectivity to the limbic network compared to non-therapeutic configurations (p=4.2×10^-3^, two-sided permutation test with Bonferroni correction).

**Figure 4.**
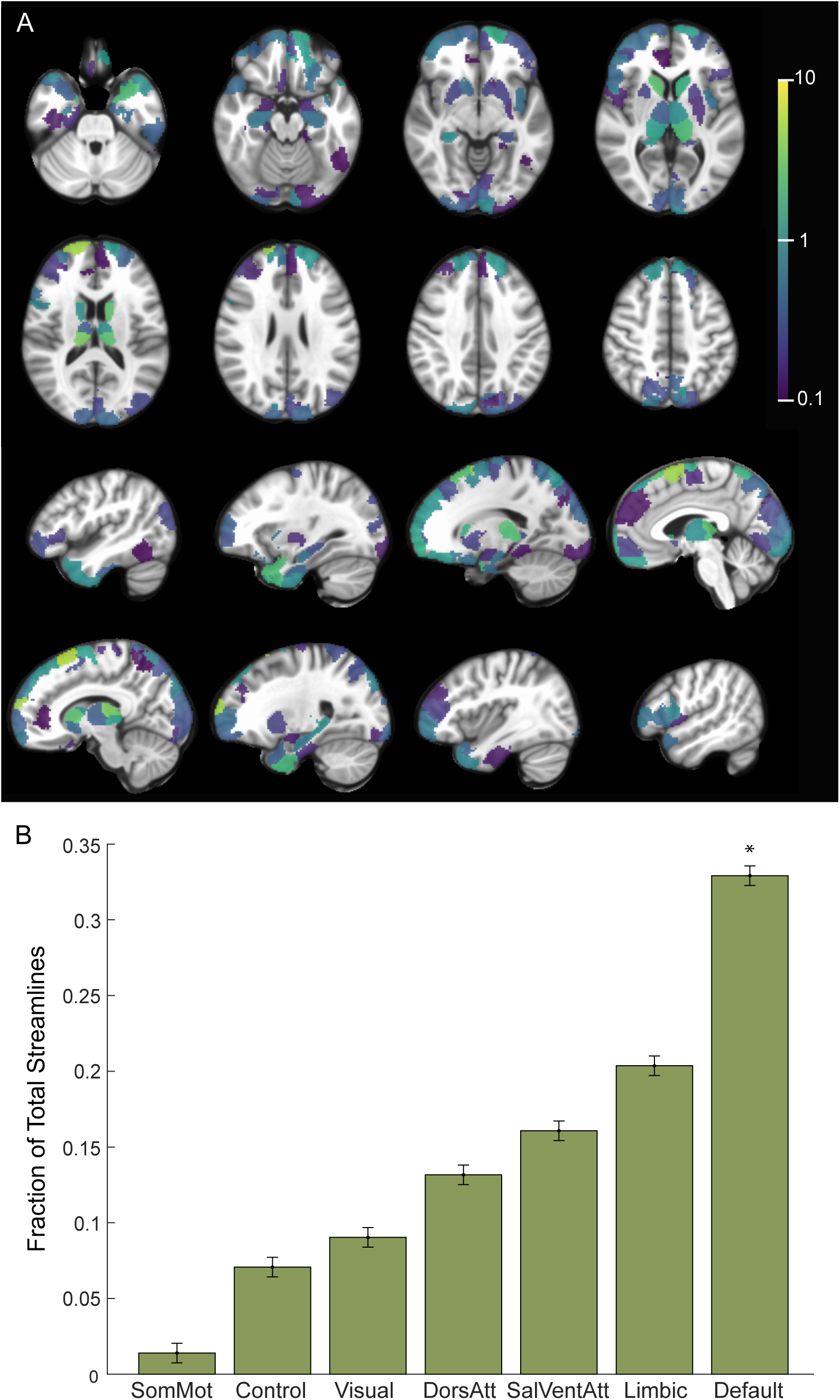
Structural Connectivity with Therapeutic Electrode Configurations. A) Percentage of total streamlines from the estimated volume of activated tissue to functional network parcels for the therapeutic configurations (n=6 runs, 3 subjects). B) Resting state networks ordered based on increasing fraction of streamline counts from the estimated volume of activated tissue for the therapeutic configurations. * p<0.05, permutation test (one-sided).

## Discussion

In this study, we developed an fMRI paradigm in which DBS is cycled On and Off to investigate the brain network mechanisms underlying this treatment. We found that DBS from therapeutic contacts induced long-range BOLD suppression in regions implicated in OCD such as the lateral orbitofrontal cortex, dorsomedial prefrontal cortex, and subthalamic nuclei. Many of these suppressions were found to be concentrated within the default mode network. In contrast, DBS configurations that were non-therapeutic often led to heterogenous and non-specific brain activation patterns. Moreover, we found that the estimated VAT nearby therapeutic DBS contacts showed significant structural connected to the DMN, but not to other networks.

Based upon these findings, we propose a model in which therapeutic ALIC DBS operates by interrupting communication through white matter to structurally connected regions such as the medial prefrontal cortex, lateral orbitofrontal cortex, thalamus, and midbrain, which are part of a CSTC circuit implicated in OCD. The suppression of these directly connected regions in turn leads to a much wider cascading set of suppressions throughout the wider DMN including regions that do not appear have direct structural connections to the leads, presumably by way of polysynaptic connections. The DMN is often associated in the literature with internalizing states [37], and prior studies have implicated disruptions in functional connectivity within the default mode network [38] and between the DMN and other networks in the pathophysiology of OCD [39]. We speculate that the suppression of DMN activity might mediate the therapeutic effects of ALIC DBS by reducing obsessions and other internalizing states.

Many components of the CSTC circuit suppressed by DBS On-vs-Off are neuromodulation targets for treating OCD. For example, we observe DBS suppression of dorsomedial PFC [40] and orbitofrontal cortex [41, 42], which are TMS targets in OCD. Similarly, the suppression observed in subcortical structures such as the subthalamic nucleus [43] are of interest given that this region is also an alternative DBS target for treating OCD. This pattern of functional suppression also mirrors structural connections whose stimulation has been associated with improved DBS outcomes for OCD [9, 11, 44, 45].

By contrast, we observe that non-therapeutic configurations fail to exert the same disruptive effect on the default mode network and its associated subcortically connected circuits. In contrast, many of the non-therapeutic stimulation configurations instead appear to enhance, rather than suppress, brain-wide activity. The fact that similar DBS parameters can have opposing functional effects within and across distributed circuits is striking. These findings are not easily reconciled with existing models which posit that DBS stimulation always behaves like a functional lesion within a restricted anatomical region or structurally connected circuit. Instead, they suggest that the impact of DBS on distributed circuits may be activating or suppressing depending on a wide variety of factors, including network state and other patient-specific factors. Indeed, a study in a rodent model of DBS for OCD described similar competing neural populations in response to ALIC DBS [46]. Our findings are also consistent with results in Parkinson’s disease, demonstrating that therapeutic STN DBS appeared to decrease BOLD activity within a motor network whereas non-therapeutic DBS seemed to recruit non-specific activity in other non-motor regions [47].

There are several limitations to this study. First, the sample includes only a small number of patients. However, by leveraging the fact that multiple DBS configurations could be trialed and compared within each subject we were nevertheless able to identify changes in network activity that were specific to therapeutic DBS. We also utilized a block design, which can yield a larger effect size than alternative study designs. Another limitation is that we could not consistently observe the local impact of DBS using fMRI due to the presence of the electrode artifact around the stimulation contacts. Lastly, it is unclear how the acute functional changes that we observe using our DBS On/Off cycling protocol are related to the long-term OCD benefit of continuous stimulation. It is possible that the suppression of the DMN with cycling DBS On and Off may be more related to acute mood and anxiety changes, rather than changes specific to the core pathophysiology of OCD [48]. Indeed, during testing for tolerability, the DBS programming clinician noted that the therapeutic DBS in our responders was uniformly associated with acute improvements in mood and reductions in anxiety. In contrast, many of the non-therapeutic settings were associated with worse anxiety following stimulation.

Our study suggests that therapeutic DBS suppresses the CSTC circuit and DMN in responders. Future studies will be needed to determine if the same acute network changes that we observe with therapeutic ALIC DBS can also be observed with other evidence-based DBS targets for OCD, such as the anteromedial subthalamic nuclei and how they relate to proposed anatomic sweet-spots for OCD DBS within the ALIC region [49]. It also remains to be seen whether the use of imaging-based biomarkers can help guide and simplify the process of DBS programming. Our results may also inform other closed-loop approaches targeted to suppress CSTC circuit activity to improve outcomes for patients with severe, refractory OCD [50].

## Acknowledgments and Disclosures

This work is supported by the National Institute of Mental Health (Grant No. K23MH125018, Grant No. R21MH130914), UCSF Team Science Research Allocation Program Grant, Foundation for OCD Research, and Foundation for the NIMH. The authors would like to thank the MRI safety personnel at UCSF for their involvement in the post-operative imaging studies. We also would like to that Lee Reid and Audrey Kist for their helpful comments on this manuscript. All authors report no biomedical financial interests or potential conflicts of interest.

## Author Contributions

Conception and design of the work, A.M.L and M.A.M; Acquisition of data A.M.L, N.S, M.A.M, A.C.F; Analysis and interpretation of the data N.S, M.A.M, A.M.L, G.B; Writing of manuscript A.M.L, N.S., M.A.M; Resources: A.M.L and M.A.M; Supervision A.M.L and M.A.M

## Data Availability

The human patient data relevant to this study are accessible under restricted access according to our IRB protocol. The de-identified patient data that support the findings of this study will be made available from the corresponding author upon request.

## Notes

### Competing Interest Statement

The authors have declared no competing interest.

