## Supplementals for "Therapeutic DBS for OCD Suppresses the Default Mode Network"

**Supplemental Table 1: DBS Response of Participants**

| **Subjects** | **Baseline YBOCS** | **Last YBOCS** | **% Change YBOCs** | **Response?** | **DBS Treatment Configuration** |
| --- | --- | --- | --- | --- | --- |
| OCD01 | 35 | 20 | 42.9% | Responder | LC+2- (5mA), RC+1- (5mA) |
| OCD_A | 36 | 21 | 41.7% | Responder | L0+1- (6.8mA), R0+1- (6.8mA) |
| OCD_B | 34 | 22 | 35.3% | Responder | L2+1- (7mA); R0+1- (5.4mA) |
| OCD02 | 39 | 38 | 2.6% | Non-Responder | Off |
| OCD03 | 30 | 30 | 0.0% | Non-Responder | Off |

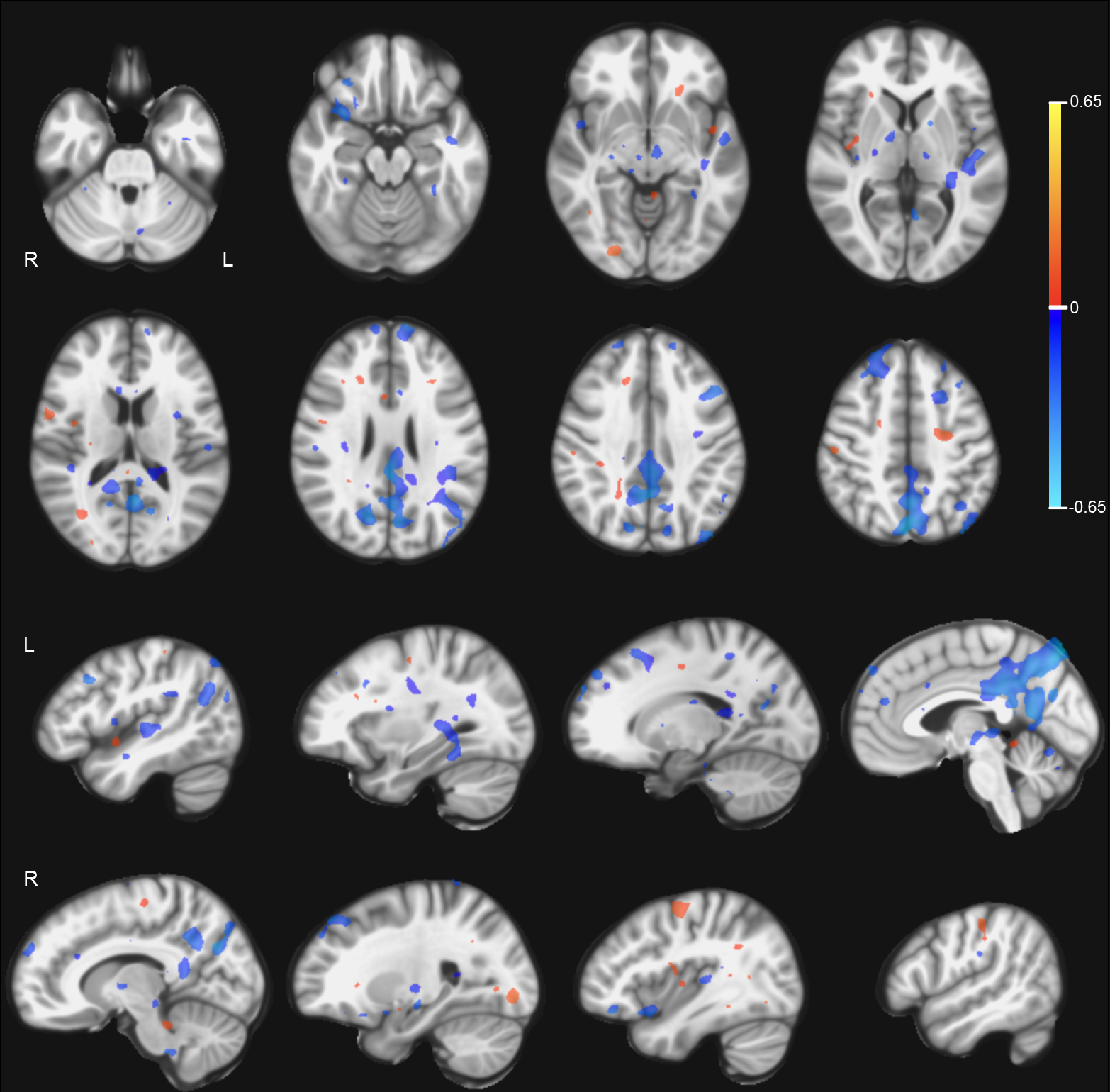

**Supplemental Fig 1. BOLD response to therapeutic DBS On vs Off.** Group comparison using linear mixed effects model of BOLD response between DBS ON vs Off in therapeutic DBS configurations. Activations are in red and suppressions in blue. Color bar indicates percentage change in BOLD signal.

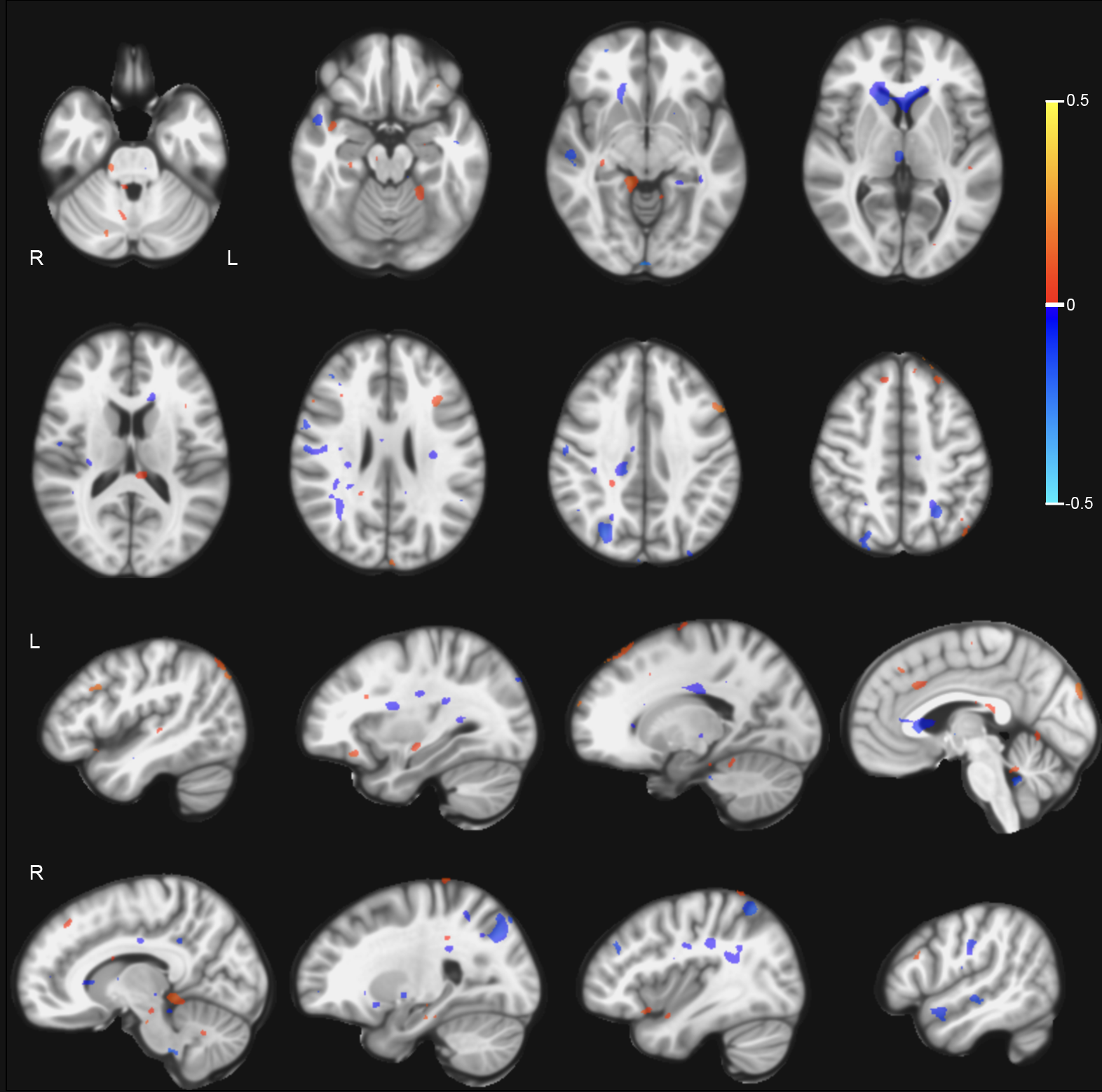

**Supplemental Fig 2. BOLD response to non-therapeutic DBS On vs Off.** Group comparison using linear mixed effects model of BOLD response between DBS On vs Off in non-therapeutic DBS configurations. Activations are in red and suppressions in blue. Color bar indicates percentage change in BOLD signal.

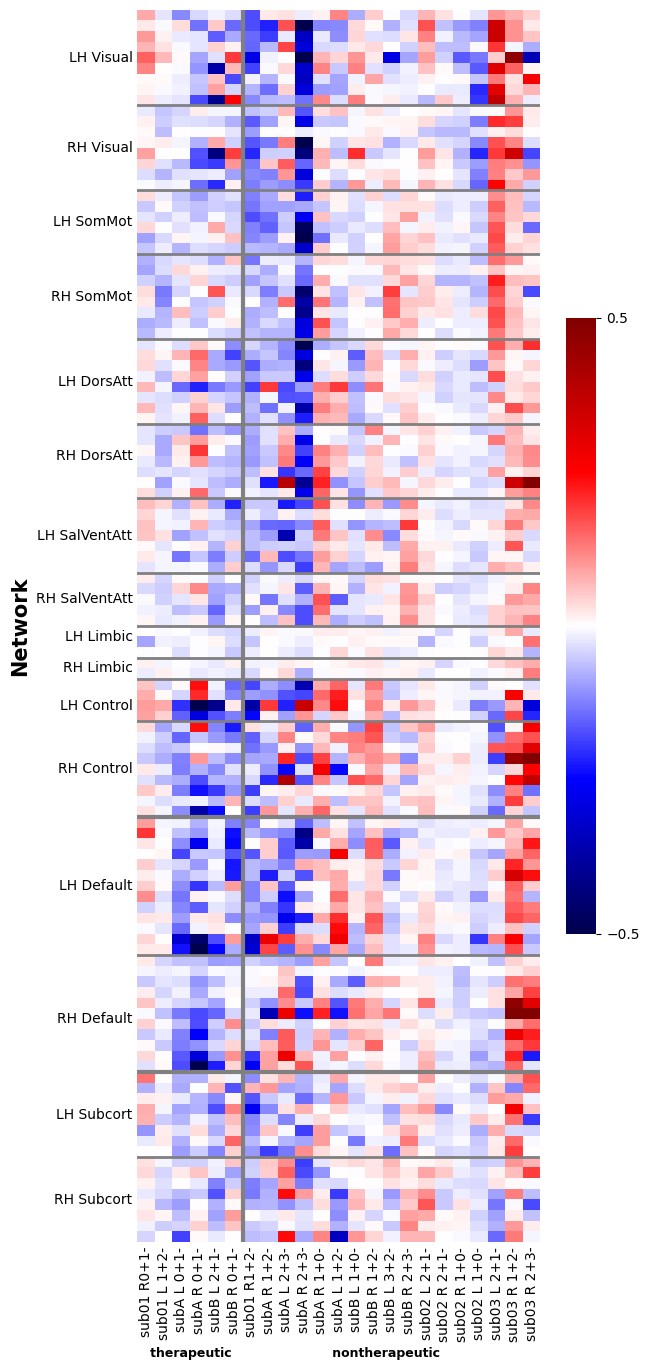

**Supplemental Fig 3. BOLD response to non-therapeutic DBS On vs Off.** BOLD response comparing across therapeutic and non-therapeutic configurations within parcels across canonical networks. Activations are in red and suppressions in blue.

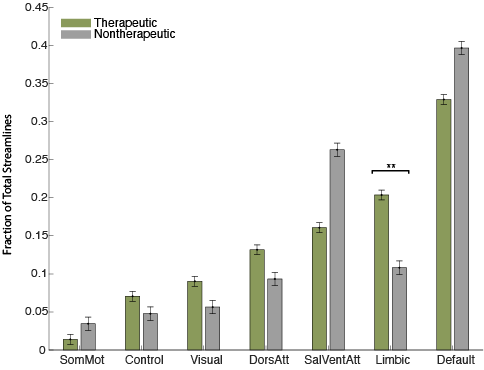

**Supplemental Fig 4. Structural Connectivity with Therapeutic and Non-Therapeutic Electrode Configurations**. A) Fraction of total streamlines from the estimated volume of activated tissue to functional network parcels for the therapeutic and non-therapeutic configurations. Cortical parcels from Schaefer 100 parcellation and subcortical structures from Melbourne subcortical atlas S1. ** p=0.005, permutation test (two-sided).
